# The matrisome of the murine and human dorsal root ganglion: a transcriptomal approach

**DOI:** 10.1101/2022.10.22.513341

**Authors:** Robin Vroman, Rahel Hunter, Matthew J. Wood, Olivia C. Davis, Zoë Malfait, Dale S. George, Dongjun Ren, Diana Tavares-Ferreira, Theodore J. Price, Anne-Marie Malfait, Fransiska Malfait, Rachel E. Miller, Delfien Syx

## Abstract

The extracellular matrix (ECM) is a dynamic structure composed of a large number of molecules that can be divided into six different categories and are collectively called the matrisome. The ECM plays pivotal roles in physiological and pathological processes in many tissues, including the nervous system. Intriguingly, alterations in ECM molecules/pathways are often associated with painful human conditions and murine experimental pain models. Nevertheless, mechanistic insight into the interplay of normal or defective ECM and pain is largely lacking. To expand the knowledge on ECM composition and synthesis in the peripheral nervous system, we used a transcriptomal approach to investigate the expression and cellular origin of matrisome genes in murine and human dorsal root ganglia (DRG), containing the cell bodies of sensory neurons. Bulk RNA sequencing data showed that over 60% of all matrisome genes were expressed in both murine and human DRG, with proportionally more core matrisome genes (glycoproteins, collagens, and proteoglycans) expressed compared to matrisome-associated genes (ECM-affiliated genes, ECM regulators and secreted factors). Examination of the cellular origin of matrisome expression by single cell RNA sequencing on murine DRG revealed that core matrisome genes, especially collagens, were expressed by vascular leptomeningeal-like (fibroblast) cell types whereas matrisome-associated genes were mainly expressed by neuronal cell types. We analyzed cell-cell communication networks with the CellChat R package and predicted an important role for the Collagen signaling pathway in connecting vascular cell types and nociceptors in murine tissue, which we confirmed by analysis of spatial transcriptomic data from human DRG. RNAscope *in situ* hybridization and immunohistochemistry confirmed expression of collagens in fibroblasts surrounding nociceptors in human DRG. This study supports the idea that the DRG matrisome may contribute to neuronal signaling in both mouse and human. The identification of the cellular distribution of murine and human matrisome genes provides a framework to study the role of the ECM in peripheral nervous tissue and its effects on pain signaling.

**Highlights:**

- Transcriptomal analyses of mouse and human dorsal root ganglia (DRG) revealed that over 60% of matrisome genes are expressed by murine and human dorsal root ganglia (DRG), with over 85% of the genes with orthologues overlapping between both species.
- Matrisome-associated genes had the highest expression in both species and included conserved expression of annexins, S100 calcium binding proteins and cathepsins.
- Collagens and collagen receptors are expressed by distinct cell types in murine and human DRG, suggesting that the collagen signaling pathway could be involved in cell-cell signaling.

## Introduction

Chronic pain is a common worldwide problem with inadequate treatment options [1,2]. Intriguingly, many pathological conditions associated with extracellular matrix (ECM) alterations are associated with the presence of chronic pain [3–7]. Indeed, pain is often the primary reason patients seek medical attention for complex diseases such as osteoarthritis, and for heritable connective tissue disorders like Ehlers-Danlos Syndromes, Marfan Syndrome, and osteogenesis imperfecta [8–12]. Nociception lies at the basis of pain perception. Nociceptors innervating peripheral tissues are activated by a painful stimulus. The generated pain signal gets transduced to dorsal root ganglia (DRG), which are a part of the peripheral nervous system and contain the cell bodies of the sensory neurons [13,14]. From the DRG, the pain signal is propagated to the spinal cord and brain, where the signal is consciously perceived as pain. The development of chronic pain involves changes at all levels of the nervous system that modify how an acute transient pain signal is processed and transforms into persistent pain. In the DRG, these pain-associated alterations can include changes in transcription patterns as well as an influx of immune cells [15–17].

The ECM is a dynamic and interactive three-dimensional network consisting of a large variety of macromolecules that provides structural support and mechanical properties to cells and tissues, including the nervous system [18–22]. Although the ECM constituents are fundamentally the same, all tissues have a unique ECM composition and topology, adapted to meet their functional requirements [23–25]. Healthy ECM is crucial for optimal functioning of the nervous system. Studies have shown that integrins and laminins are important for neurite outgrowth from DRG neurons *in vitro*, and that ECM-integrin signaling are crucial for regenerative axon assembly [26–28]. The ECM also provides stability for the laminin dependent myelination of the peripheral nerves [29].

An exact overview of which ECM genes are expressed in DRG tissue is lacking, which prohibits further understanding of the roles the ECM plays in nociceptive functioning. Naba *et al*. created a list of ‘matrisome’ genes as an ECM framework, which contains structural core matrisome genes, such as glycoproteins, collagens, or proteoglycans, as well as matrisome-associated genes, including signaling molecules and enzymes [30,31]. In this study, we investigated which matrisome genes were expressed in human and murine DRG, described the cellular distribution of the expressed matrisome genes in the DRG, and investigated potential interactions between cell types in the DRG through ligand-receptor analyses, RNAscope *in situ* hybridization and immunohistochemistry (IHC).

## Results

### Similar percentages of matrisome genes are expressed by murine and human DRG

To obtain an overview of the expression of matrisome genes in DRG, bulk RNAseq was performed on murine and human DRG and expressed genes were filtered against publicly available lists of murine (n = 1,110) and human (n = 1,027) matrisome genes [32].

In murine DRG collected from lumbar levels L3-L5, 65±0.35% of the 1,110 murine matrisome genes were expressed (transcripts per million (TPM) > 0.1; male: n = 6, female: n = 5). Of the 274 core matrisome genes, 83±0.38% were expressed, which was significantly higher compared to 59±0.42% of the 836 matrisome-associated genes (p < 0.0001) (Supplemental Figure S1A, Supplemental Table 1). Specifically, for the core matrisome, 81±0.38% of glycoprotein, 88±0.55% of collagen, and 86±1.1% of proteoglycan genes were expressed; whereas for the matrisome-associated genes 69±0.61% of extracellular matrix (ECM)-affiliated, 56±0.36% of ECM regulators and 58±0.58% of secreted factors genes showed expression (Figure 1A). We did not see any sex specific differences in overall expression levels in murine DRG (Supplemental Figure S1A).

**Figure 1:**
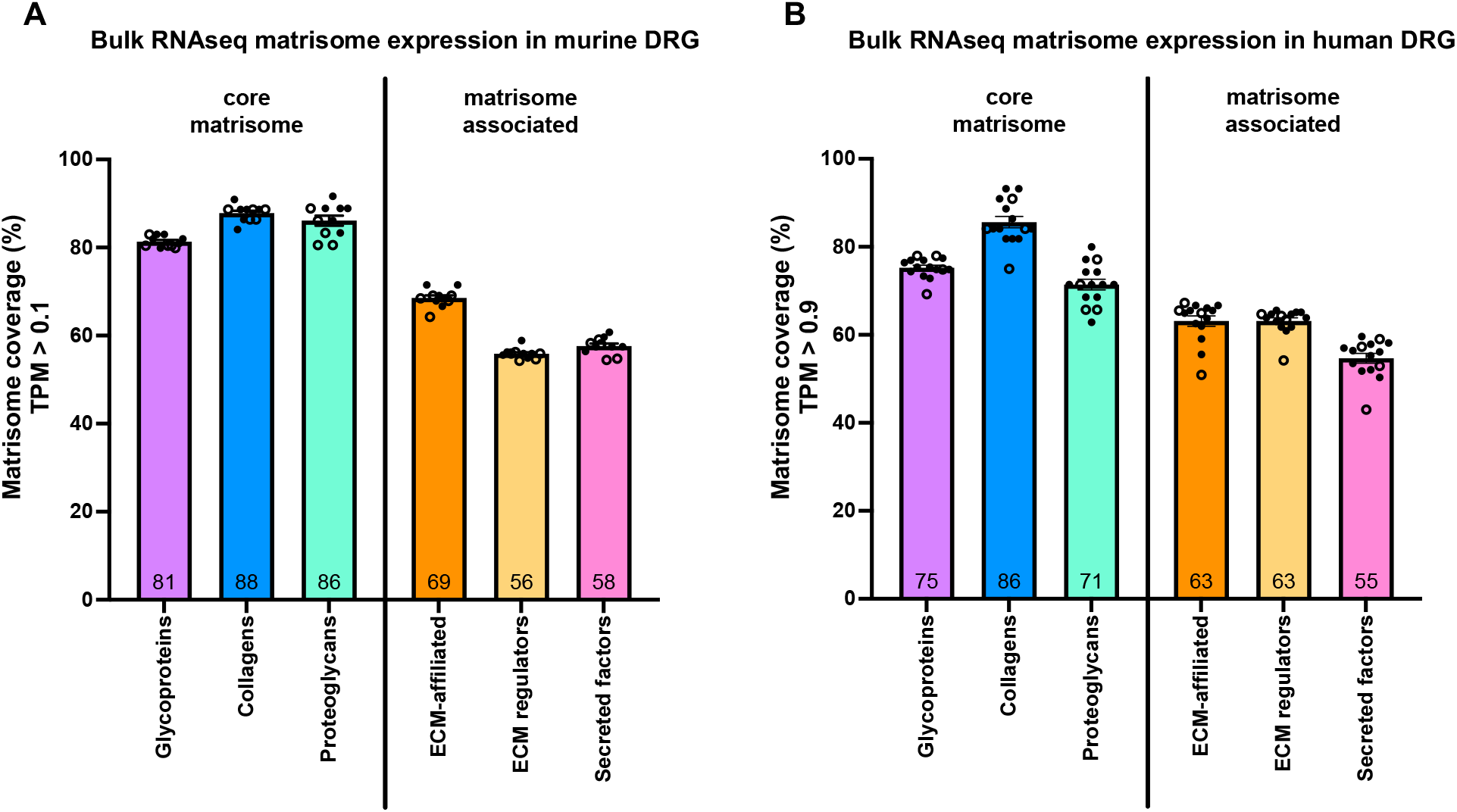
Matrisome gene expression in murine and human DRG. **A)** Bulk RNAseq was used to identify the percentage of each category of matrisome genes expressed in murine DRG. Dots represent the percentage of expressed genes in DRGs collected from one mouse. n = 6 male (filled dots), n = 5 female (open dots). Total number of genes per category: 194 glycoproteins, 44 collagens, 36 proteoglycans, 165 ECM-affiliated proteins, 304 ECM regulators, 367 secreted factors. **B)** Bulk RNAseq was used to identify the percentage of each category of matrisome genes expressed in human DRG. Dots represent the percentage of expressed genes in DRGs collected from one individual. n = 11 male (filled dots), n = 4 female (open dots). Total number of genes per category: 195 glycoproteins, 44 collagens, 35 proteoglycans, 171 ECM-affiliated proteins, 238 ECM regulators, 344 secreted factors. Number inside the bar represents the mean per group. Mean±SEM. Human bulk RNAseq data was previously published [33].

When interrogating previously published bulk RNAseq data from human DRG for human matrisome genes, a similar trend was observed [33]. Overall, 64±0.87% of the 1,027 human matrisome genes were expressed (TPM > 0.9; male: n = 11, female: n = 4). Of the 274 core matrisome genes, 76±0.63% were expressed, which was significantly higher than 59±0.97% of the 753 matrisome-associated genes (p < 0.0001) (Supplemental Figure S1B, Supplemental Table 2). In more detail, for the core matrisome we found that 75±0.59% of glycoproteins, 86±1.3% of collagens, and 71±1.2% of proteoglycans were expressed (Figure 1B). For the matrisome-associated genes 63±1.2% of ECM-affiliated, 63±0.73% of ECM regulators, and 55±1.1% of secreted factors were expressed above the threshold (Figure 1B). Furthermore, no sex-specific differences were seen in the ratios of matrisome genes being expressed between male and female human DRG (Supplemental Figure S1B).

Next, we compared the expressed genes in order to determine whether or not the same genes were being expressed in mouse and human. Starting with the human matrisome list, we filtered out all genes being expressed in the human DRG that have at least one murine ortholog (660 genes) and found that 86% of these genes were also being expressed in the murine DRG (592 genes).

These analyses demonstrated that a substantial proportion of the *in silico* defined matrisome genes are expressed in both murine and human DRG and in similar levels in both male and female, however there are some differences in the exact genes expressed in each species.

### Highest expressed matrisome genes in murine and human DRG

To examine the highest expressed matrisome genes in murine and human DRG, matrisome genes were ranked based on their average TPM values and analyzed either across all matrisome categories or per matrisome category separately (Figure 2, Supplemental Figure S2, Supplemental Table 1-2).

**Figure 2:**
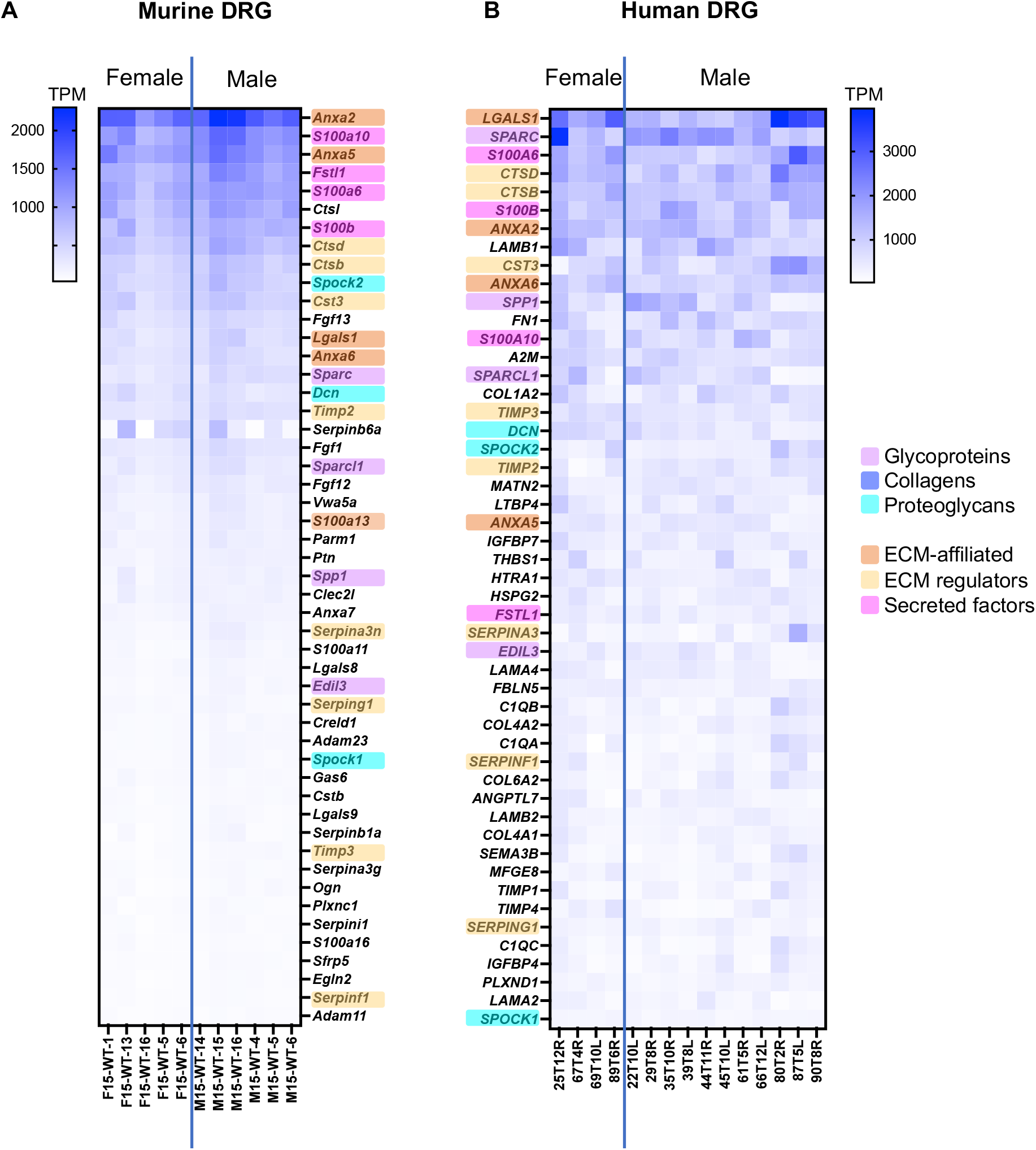
Bulk RNAseq was used to identify the 50 highest expressed matrisome genes in murine and human DRG. **A)** Murine matrisome genes are ranked by average TPM value across all samples. Male: n = 6, female: n = 5. **B)** Human matrisome genes are ranked by average TPM value across all samples. Male: n = 11, female: n = 4. Orthologues between murine and human datasets have been highlighted in the color corresponding to the matrisome categories. Human bulk RNAseq data was previously published [33].

Across all matrisome categories, we focused on the 50 genes with the highest TPM value, which corresponds to approximately the highest 5% of all matrisome genes. For murine DRG, 11 out of the 50 highest expressed genes were core matrisome genes compared to 23 out of 50 for the human DRG. Between mouse and human, 23 matrisome genes overlapped in both 50 highest expressed gene lists (46%) (Figure 2). Seven of these 23 overlapping genes were core matrisome genes (4 glycoproteins and 3 proteoglycans), while 16 overlapping genes belonged in the matrisome-associated group (4 ECM-affiliated genes, 8 ECM regulators, and 4 secreted factors). In particular, matrisome-associated genes had the highest expression in both species and included conserved expression of annexins (*Anxa2, Anxa5*, and *Anxa6*), S100 calcium binding proteins (*S100a6, S100a10* and *S100b*), and cathepsins (*Ctsb, Ctsd*, and *Cts3*) (Figure 2).

Among the non-overlapping genes between species, some key differences in expression levels were observed, which may be related to the overall difference in cellular content between mouse and human [14]. For example, more collagen (*COL1A2, COL4A1, COL4A2*, and *COL6A2*) and laminin (*LAMA2, LAMA4, LAMB1* and *LAMB2*) genes made the 50 highest expressed gene list for human compared to mouse, which may reflect the fact that human DRG have more fibrous content than mouse DRG. In contrast, mice had more fibroblast growth factor (*Fgf1, Fgf12*, and *Fgf13*) genes in the 50 highest expressed genes list, consistent with their expression by DRG neurons in other published murine data sets [34]. To look in more detail in each matrisome category, we also compiled lists of the 10 highest expressed genes in each category (Supplemental Figure S2). From these lists we can see again that, while many of these genes are conserved between species, the relative expression levels differ in murine and human DRG – this can be particularly noted in the ‘collagens’ category.

Examination of the heatmap resulting from ranking the matrisome genes according to expression levels revealed less intersample variability per gene in the mouse dataset compared to the human dataset (Figure 2). The smaller variability observed in the mouse samples compared to the human samples was expected based on a more variable cohort of human samples compared to age-matched inbred mice.

Despite the fact that the murine and human matrisome gene ensembles are not identical, we observed overlap in the overall and category-specific highest expressed matrisome genes between murine and human DRG.

### Cellular distribution of matrisome gene expression in murine DRG

To pinpoint the cellular origin of matrisome gene expression and to elucidate which DRG-resident cells express these genes, single cell RNA sequencing (scRNAseq) was performed on murine DRG (L3-L5 unilateral, pooled from 10 male mice, 18 weeks of age). A total of 8,755 cells were clustered into different cell types. Based on cluster specific markers, eight different cell types were identified, including neuronal cells (nociceptors (NOCI) and large diameter neurons (LDN)), supporting cells (Schwann cells (SCHW) and satellite glial cells (SATG)), vascular cell types (vascular leptomeningeal cells/fibroblast-like cells (VLMC-like), vascular endothelial cells (VEC), and vascular smooth muscle cells arterial (VSMCA)), and immune cells (IMM) (Supplemental Figure S3A-B) [35,36]. As a validation, gene expression for each matrisome category was checked in the scRNAseq data and percentages of expressed genes were overall consistent with the bulk RNAseq results, indicating that the obtained scRNAseq dataset is representative for subsequent matrisome analysis (Supplemental Figure S3C).

To investigate the cellular origin of the matrisome genes, we focused on the 25 highest expressed genes per category obtained from the murine bulk RNAseq data and checked their expression in the different cell clusters from the scRNAseq dataset (Figure 3, Supplemental Table 3). For the core matrisome, cell type-specific expression patterns were observed. Glycoproteins were expressed by all cell types in the DRG, with the exception of immune cells. Although each glycoprotein gene had its own cellular distribution pattern, overall, VLMC-like cells (fibroblasts) were expressing most of the 25 highest expressed glycoproteins. Collagens were predominantly expressed by vascular cell types, more specifically VLMC-like cells, and not by neuronal or immune cell types. Finally, proteoglycans were primarily expressed by vascular cell types and to a lesser degree by neuronal cells. Two genes deviated from this pattern, *Ogn* and *Srgn*, encoding the proteoglycans encoding osteoglycin and serglycin, respectively. These genes were expressed by Schwann cells and immune cells, respectively, which is consistent with the literature [34,35,37].

**Figure 3:**
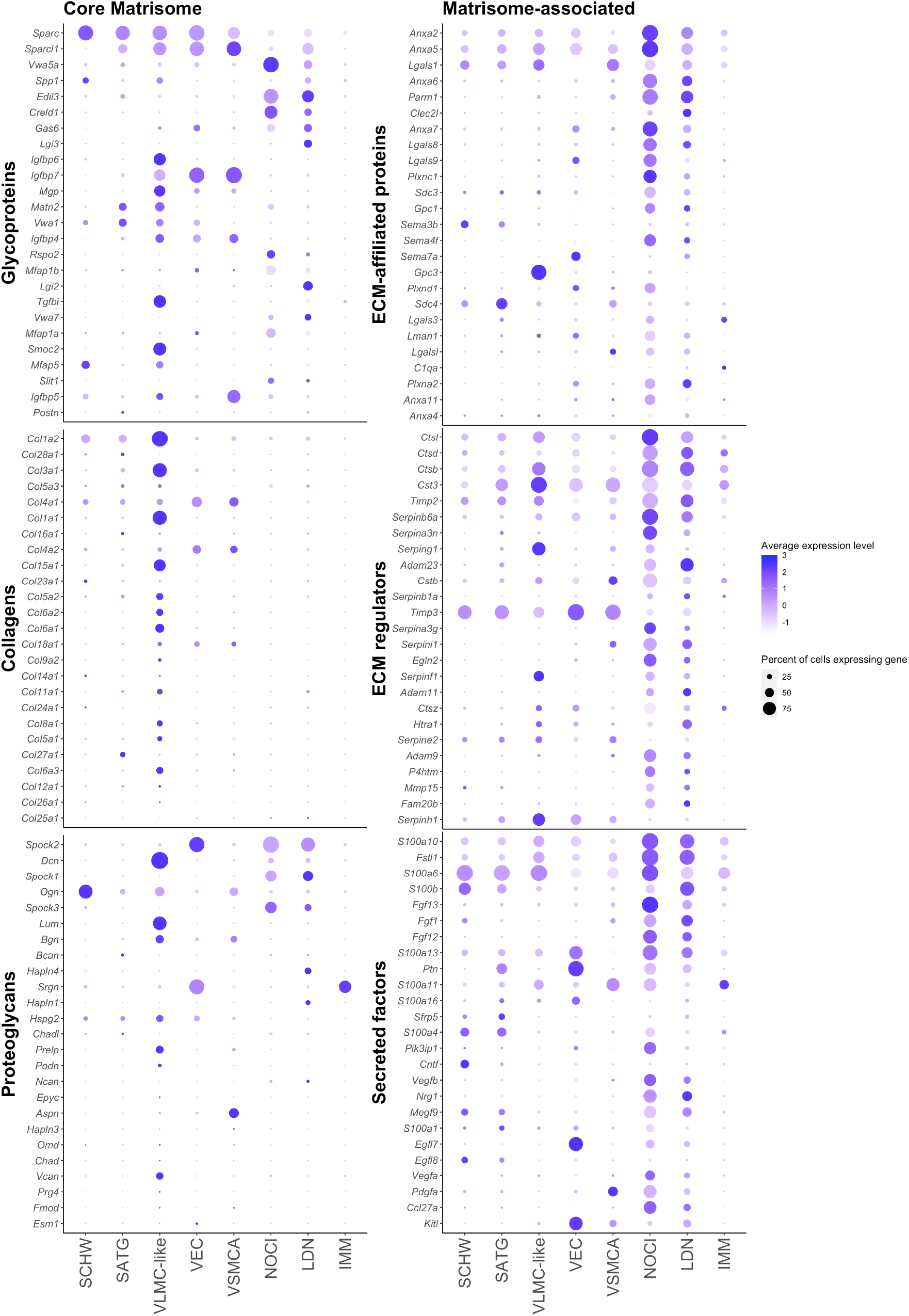
Cellular distribution of the 25 highest expressed genes of each matrisome category in murine DRG determined by scRNAseq. The size of each dot represents the percentage of cells expressing the given gene within a cluster, and the color of each dot corresponds to the average expression (scaled data) across all cells within a cluster for each gene of interest. Schwann cells (SCHW), satellite glial cells (SATG), vascular leptomeningeal like cells (VLMC-like), vascular endothelial cells (VEC), vascular smooth muscle cells arterial (VSMCA), nociceptors (NOCI), large diameter neurons (LDN) and immune cells (IMM).

The expression of matrisome-associated genes was predominantly driven by neuronal cell types, with all three categories being expressed by both nociceptors and large diameter neurons (Figure 3). Other DRG-resident cell types also expressed matrisome-associated genes but in general with a lower expression level and by a lower percentage of cells.

### Matrisome ligand-receptor interactions in the DRG

To further elucidate the role of the ECM in DRG, interactions between different cell types within the DRG were examined based on the murine scRNAseq data and by using the R package CellChat (v1.1.3). CellChat is an integrated cell-cell communication tool that examines scRNAseq datasets and predicts cell-cell interactions and infers involved pathway networks [38].

After analyzing the full set of murine DRG scRNAseq data with CellChat, including both matrisome and non-matrisome genes, we found that all cell types had the potential to interact with each other with various predicted strengths (Supplemental Figure S4A-B). Subsequently, a list of enriched ligand-receptor pathways was generated (Supplemental Figure S4C). Interestingly, several pathways correlating with the constituents of the core matrisome were found to be highly enriched, such as the ‘Collagen’, ‘Fn1’, and ‘Hspg’ pathways. In addition, pathways related to the matrisome-associated genes, were also found to be enriched, including the ‘Fgf’, ‘Sema3’, and ‘Vegf’ pathways. The Collagen signaling pathway was the highest ranked and as such contributed the most to the predicted interactions occurring in the DRG (Figure 4A-B, Supplemental Figure S4). In particular, *Col1a1, Col1a2, Col4a1* and *Col4a2*, encoding types I and IV collagen, were identified as ligands that can interact with the receptors *Cd44* and *Sdc4*, encoding cluster of differentiation 44 and syndecan-4, respectively. Other receptors contributing to collagen interactions included the integrin pair consisting of *Itga1* and *Itgb1*, which encode the α1 and β1 subunits (Figure 4B). Interactions with less weight involved the type VI collagen encoding genes, *Col6a1, Col6a2* and *Col6a3*, as ligands, and *Sdc1*, encoding syndecan-1, and the integrins *Itga3, Itgav*, and *Itgb8*, encoding the α3, αv, and β8 subunits, as receptors (Figure 4B). Examining our scRNAseq data allowed us to examine the cellular sources of these genes in more detail. This revealed that the ligands of the Collagen pathway were primarily expressed by the vascular cell types, in particular the VLMC-like cells (fibroblasts), and to a lesser extent by the supporting Schwann cells and satellite glial cells (Figure 4C, Supplemental Figure S5A). The cell types expressing the interacting receptors were more diverse. *Sdc4* was expressed by satellite glial cells, Schwann cells, VLMC-like and VSMCA cells, whereas *Cd44* was expressed by nociceptors and immune cells. The integrin pair *Itga1* and *Itgb1* was expressed by all other cell types in murine DRG (Figure 4D, Supplemental Figure S5B). Large diameter neurons did not express any of these ligands or receptors at a high enough level to be included in these Collagen pathway ligand-receptor analyses.

**Figure 4:**
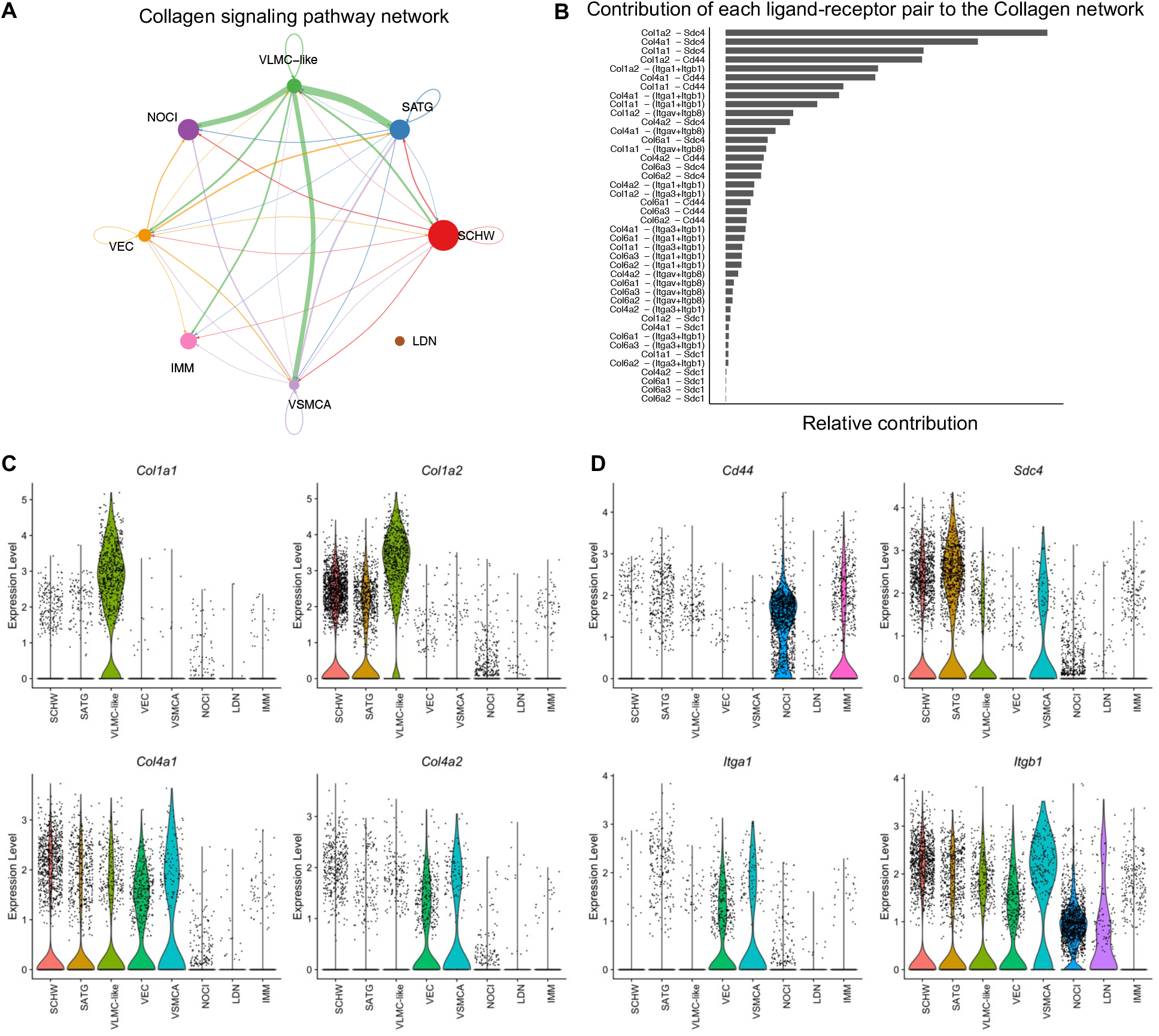
Cell-cell communication in murine DRG inferred from scRNAseq data. **A)** The inferred ‘Collagen’ signaling network in murine DRG. Arrow points show the direction of the interaction, and the size of the arrow corresponds with the weight of interaction. Size of the dots corresponds with the relative number of cells per cell cluster. Schwann cells (SCHW), satellite glial cells (SATG), vascular leptomeningeal like cells (VLMC-like), vascular endothelial cells (VEC), vascular smooth muscle cells arterial (VSMCA), nociceptors (NOCI), large diameter neurons (LDN) and immune cells (IMM). **B)** Each predicted ligand-receptor pair within the Collagen signaling network in murine DRG is ranked based on the relative contribution to the overall Collagen signaling pathway in panel A. **C)** Violin plots of the most contributing ligand genes in scRNAseq murine DRG: *Col1a1, Col1a2, Col4a1, Col4a2*. **D)** Violin plots of the most contributing receptor genes in scRNAseq murine DRG: *Cd44, Sdc4, Itga1, Itgb1*.

This data suggests that matrisome genes in the Collagen signaling pathway are involved in cell-cell interactions in murine DRG and that these interactions involve ligands and receptors expressed by different cell types. The next step was to look if this cell-type specific expression pattern is present in human DRG as well. Therefore, a previously published spatial transcriptomics dataset on human DRG (male: n = 4, female: n = 4) [39] combined with an unpublished in-house human DRG sample (female: n = 1) were examined. To focus on cells within the DRG, barcodes were selected based on the H&E images (Figure 5A). We next defined markers for specific cellular subsets in the DRG: *SCN10A* functions as a widely accepted marker for nociceptors, and *DCN* was identified as a marker for VLMC-like cells (fibroblasts) [40]. Barcode spots with expression of a gene of interest above the 25^th^ quartile were considered positive (Figure 5B, Supplemental Figure S6A), after which ratios of co-expression with *SCN10A* and *DCN* were calculated. Based on the murine CellChat data, expression of collagen genes (*COL1A1, COL1A2, COL4A1, COL4A2, COL6A1, COL6A2, COL6A3*) was evaluated and showed significantly more co-expression with *DCN* compared to *SCN10A* (Figure 5C, Supplemental Figure S6B). Subsequent examination of their predicted receptors showed significantly more co-expression of *CD44* with *SCN10A* (p < 0.0001), consistent with the murine data, and of *SDC4* with *DCN* (p = 0.0199). Integrins showed a variable pattern with *ITGB1* (p < 0.0001), *ITGA3* (p = 0.0010) and *ITGAV* (p = 0.0257) co-expressing significantly more with *SCN10A, ITGB8* co-expressing more with *DCN* (p = 0.0147), and no difference in co-expression for *ITGA1* and *SDC1* (Figure 5D, Supplemental Figure S6C).

**Figure 5:**
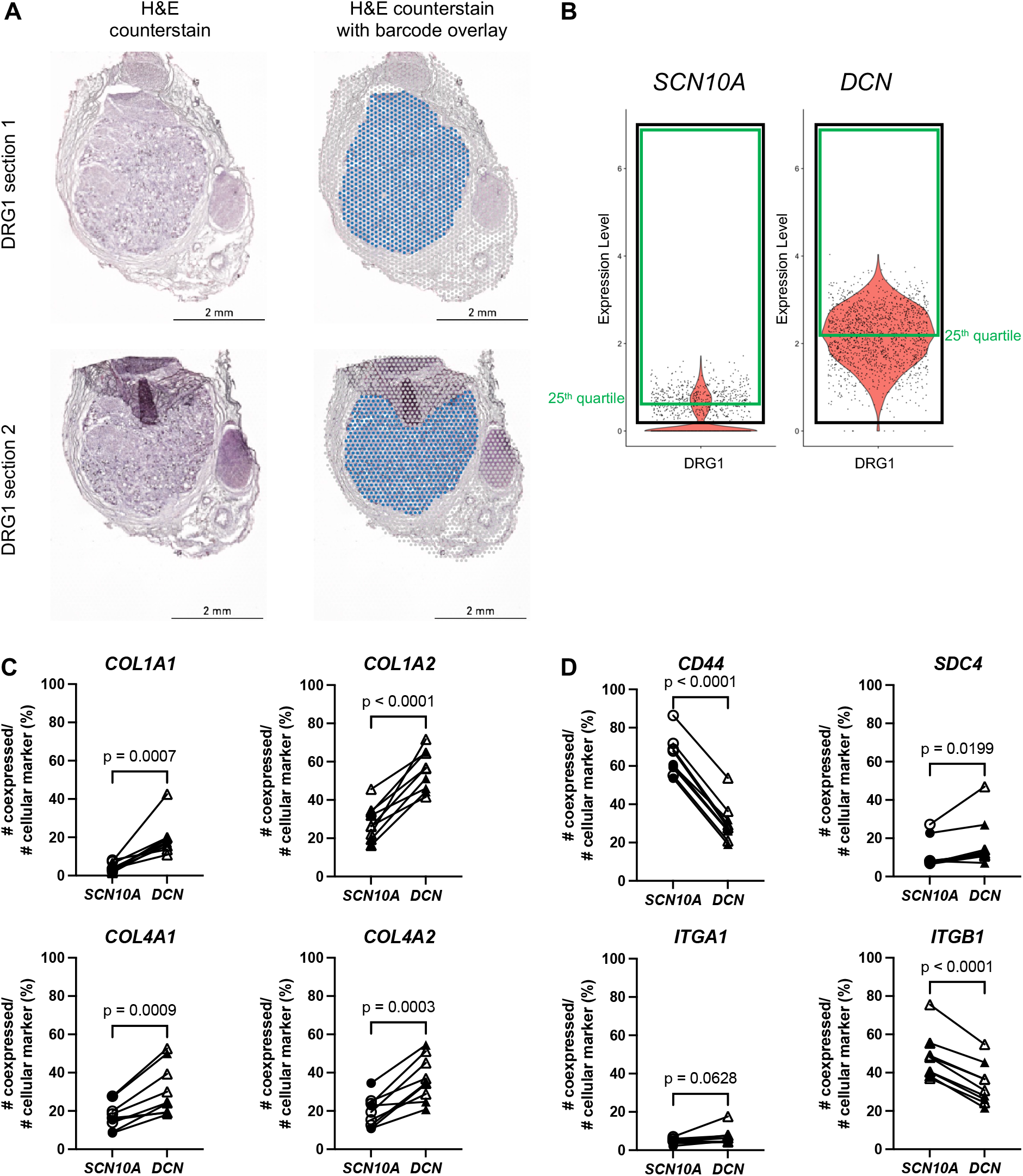
Spatial transcriptomics co-expression on human DRG. **A)** Left: H&E counterstain on two sequential sections of a human DRG (in-house sample). Right: Sections shown with the Visium barcode overlay to demonstrate which barcodes were selected for further analysis in the Seurat R package (in blue). **B)** After scaling and normalization in Seurat, expression thresholds were established for each gene of interest by using the 25th quartile cutoff (green box) of barcodes > 0 (black box) for that gene in each DRG sample. *SCN10A* and *DCN* are shown here as examples for one DRG, but this method was also applied to genes in panels C and D. **C)** Percentage of co-expression of *COL1A1, COL1A2, COL4A1*, and *COL4A2*, respectively, with *SCN10A* or *DCN*. **D)** Percentage of co-expression of *CD44, SDC4, ITGA1*, and *ITGB1*, respectively, with *SCN10A* or *DCN. SCN10A*-*DCN* double positive cells were excluded from analyses. Male: n = 4 (filled symbol), female: n = 5 (open symbol).

Overall, the human spatial transcriptomics data showed consistencies with the murine CellChat data. In particular, the examined collagen genes were preferentially expressed by VLMC-like cells (fibroblasts), and the *CD44* receptor was expressed by nociceptors. Expression of *SDC4* and the examined integrin genes was less clear-cut and is consistent with the finding that these genes were also expressed by other cell types in the murine DRG such as Schwann cells, satellite glial cells, vascular cells, and neuronal cell types.

The murine scRNAseq data indicated that collagens were barely expressed by nociceptors. To confirm this in human samples, RNAscope *in situ* hybridization was performed on human DRG sections collected from two donors with probes for *COL1A1* combined with *SCN10A*, a marker for nociceptors, or with *DCN*, a marker for VLMC-like cells (Figure 6A-F, Supplemental Figure S7). *SCN10A* positive cells did not co-express with *COL1A1*, while in 23±2.1% of DRG cells, co-expression of *COL1A1* with *DCN* was observed. We also found that 16±1.3% of *COL1A1* positive cells did not co-express with *DCN*. These findings confirm our earlier results from the murine scRNAseq data in human DRG samples. While it is valuable to know which cell types express certain genes, there can be a different dynamic at the protein level. Hence, we performed IHC on human DRG samples for the VLMC-like cell/fibroblast cell marker, platelet derived growth factor receptor α (PDGFRA), combined with the nociceptor marker transient receptor potential cation channel subfamily V member 1 (TRPV1) (Figure 6G). This demonstrated that VLMC-like cells/fibroblasts are in close proximity to TRPV1-positive cells, increasing the likelihood for the predicted interactions of collagen proteins and their receptors expressed by nociceptors or other cell types in the DRG.

**Figure 6:**
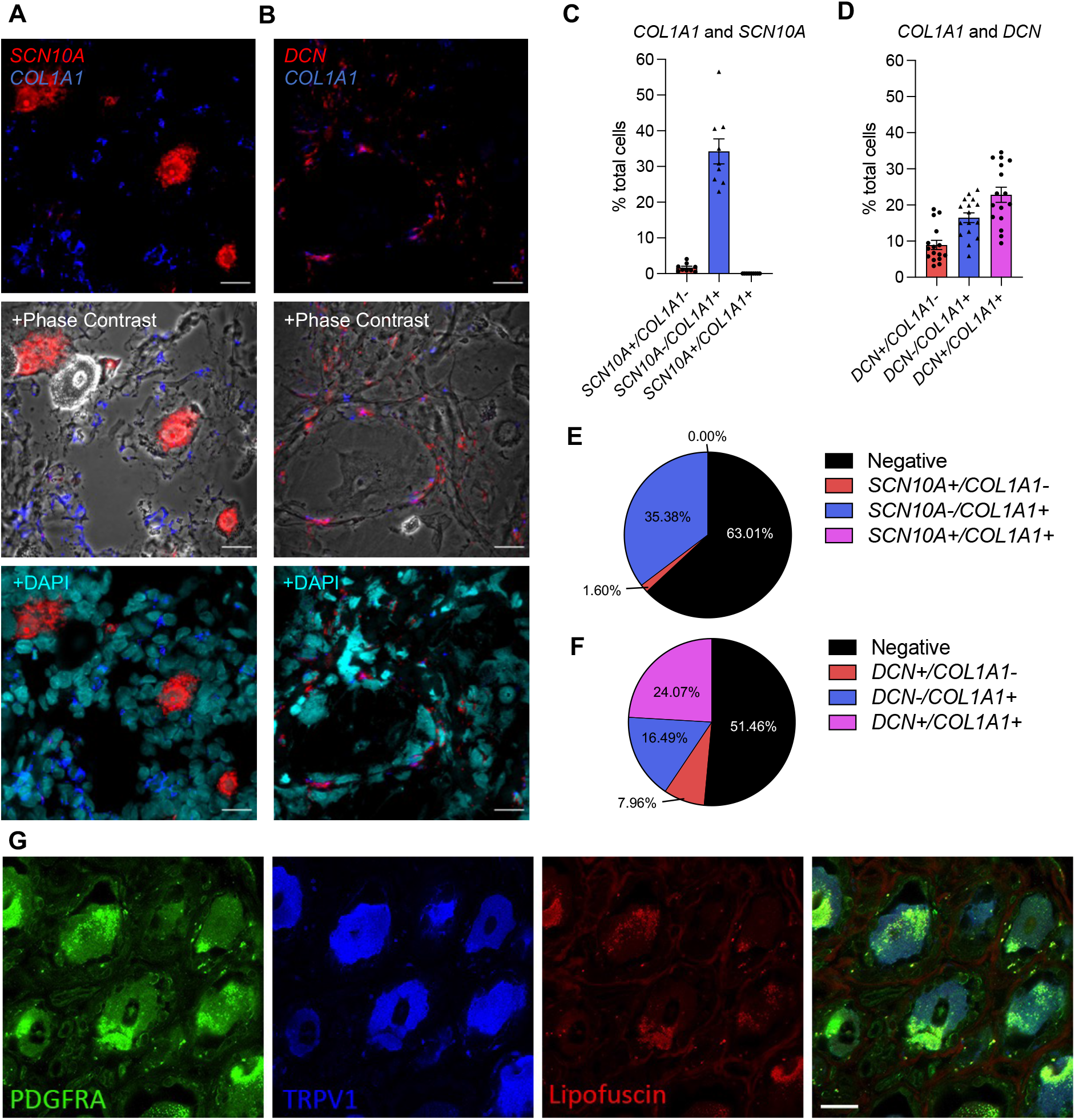
Spatial localization of collagen-producing cells in the human DRG. **A-F)** Expression of *COL1A1, SCN10A* and *DCN* in human DRG using RNAscope. **A**,**C**,**E)** RNAscope was used to identify and quantify cells expressing *SCN10A* (neuronal marker) and *COL1A1* or **B**,**D**,**F)** *DCN* (VLMC-like/fibroblast marker) and *COL1A1* in human DRG. Representative sections are shown for *COL1A1* staining with the cellular marker, either with phase contrast or nuclei staining (DAPI) overlay. (male, n=2, number of cells: *SCN10A* = 1625, *DCN* = 2359); Mean±SEM. **G)** IHC on human DRG for fibroblast marker PDGFRA, ion channel TRPV1, and the non-specific auto-fluorescent staining by lipofuscin (male: n=1, female: n=1). Each staining is represented separately and the overlay of all 3 stainings is shown. Scale bar = 25μm (A, B, G).

Taken together we found that in murine DRG, vascular cell types, including VLMC-like, VEC, and VSMCA cell, were responsible for a large portion of the expression of core matrisome genes, whereas expression of matrisome-associated genes was attributed to nociceptors and large diameter neurons. After predicting a major role for the Collagen signaling pathway in ligand-receptor interaction analyses, we showed that in human DRG, collagens were expressed by VLMC-like cells (fibroblasts) and not by nociceptors, consistent with the murine scRNAseq data. In addition, we found that VLMC-like cells (fibroblasts) are in close proximity to nociceptors, increasing the likelihood for ECM pathways to influence nociceptors and subsequent pain signaling.

## Discussion

In this study, we investigated matrisome gene expression and its cellular distribution in murine and human DRG. We found that a large percentage of matrisome genes were expressed in both murine and human DRG, with different cell types responsible for producing different types of matrisome components. Single cell and spatial transcriptomics data suggests that the ECM may contribute to cell-cell communications within the DRG, and our RNAscope *in situ* hybridization and IHC data support the idea that fibroblasts (VLMC-like cells) produce collagens and surround the neuronal cell bodies in the human DRG. These observations provide a basis for examining the functional implications of the matrisome in the DRG both in homeostasis and in chronic pain states.

Our bulk RNAseq transcriptomic experiments showed that a large fraction of matrisome genes were expressed in murine and human DRG. Murine and human matrisome lists are not identical but have a highly similar number of genes per matrisome category. The ratio of core matrisome genes being expressed was higher compared to the matrisome-associated genes for murine and human DRG, which is to be expected given the basic structural functions that glycoproteins, collagens, and proteoglycans fulfill. However, the highest expressed genes were matrisome-associated genes such as annexins (*Anxa2, Anxa5*, and *Anxa6*), S100 calcium binding proteins (*S100a6, S100a10* and *S100b*) and cathepsins (*Ctsb, Ctsd*, and *Cts3*). The annexin family has been studied in DRG for a few decades [41]. Annexins are calcium and phospholipid binding proteins and can play a role in ion channel regulation [42], including regulation of *Trpv1* [43,44] and *Trpa1* [45], ion channels important for nociception. S100 proteins have also been found in the DRG and have been shown to co-localize particularly with large diameter neurons in the DRG [46]. S100 proteins can modulate neuronal stimulation using their calcium binding properties [47], including ion channels in the DRG, by interacting with downstream receptors or mediators of K^+^ channels, Na_V_1.8 channels, or Na^+^ channels [48]. Cathepsins are lysosomal proteases that are involved in many different processes. Specifically, in a neuronal context, they have been described to be involved in neuronal development and to play a role in neurological diseases in the central nervous system [49]. In addition, a role in axon outgrowth of sensory neurons in the DRG has also been described [50]. This literature overview shows the importance of the highest expressed matrisome-associated genes in our murine and human DRG bulk RNAseq datasets, and demonstrates why it is not surprising that these were among the highest expressed genes given their involvement in neuronal functioning.

Structurally, mouse and human DRG tissue is quite different, with a higher neuronal density in murine DRG compared to human DRG, which contains thicker layers of ECM throughout the DRG [14]. Even though a lot of neurobiology has been elucidated in rodents, high quality analgesic treatments for humans remain elusive, highlighting the importance of research on human nervous tissue as well [51–53]. Here, we found that in human DRG, higher expression of collagen (*COL1A2, COL4A1, COL4A2*, and *COL6A2*) and laminin (*LAMA2, LAMA4, LAMB1*, and *LAMB2*) genes were observed compared to mice. This is consistent with the observation that human DRG have more ECM between neurons, resulting in less dense neuronal content compared to murine DRGs [14]. In contrast, mice had higher expression of genes primarily expressed by neurons such as fibroblast growth factors (*Fgf1, Fgf12*, and *Fgf13*) [34].

Recently, crosstalk between neuronal and non-neuronal cells has received increasing attention [54]. Gaining knowledge on which cells express which molecules is important for future targeted approaches trying to intervene in specific pathways. Our single cell transcriptomics data using murine DRG elucidated the cellular distribution of matrisome gene expression and demonstrated that while a large percentage of DRG nociceptors and large diameter neurons expressed the matrisome associated genes at high levels, collagens were barely expressed by neuronal cell types in mice. Knowing which cells express which molecules is the first step, but understanding the interactions between these expressed genes and different cell types is critical for elucidating potential ECM functions in the DRG. Surprisingly, when taking an unbiased approach and inputting our full list of scRNAseq data (including both matrisome genes and non-matrisome genes), we found that the Collagen pathway, a non-neuronal signaling pathway, was the most prominent ligand-receptor pathway represented in the DRG. The main interactions of this pathway involved the pro-α1 and pro-α2 chains of type I procollagen as ligands and CD44 and syndecan-4 as receptors. While the collagens were mainly expressed by VLMC-like cells (fibroblasts), CD44 was mainly expressed by nociceptors and syndecan-4 was expressed by Schwann cells, satellite glial cells, and vascular cell types. Additionally, we also examined these murine DRG interactions in a human spatial transcriptomics dataset. Genes encoding the pro-α1 and pro-α2 chains of type I collagen were also more co-expressed with a VLMC-like (fibroblast) marker, *DCN*, while the *CD44* receptor was more expressed by nociceptors. This suggests how, despite collagens being barely or not expressed by nociceptors, they could still influence nociceptive signaling.

After injury, the ECM and its associated enzymes have important functions to remove ECM debris from the injury site as well as ECM remodeling to heal the tissue properly. Also, collagens, laminin, or fibronectin are crucial in peripheral nervous tissue regeneration, and enzymes such as tissue type plasminogen activator are also important for remodeling of the ECM network [55,56]. Therefore, one can hypothesize that altered ECM homeostasis might have a detrimental effect on nociceptor functioning, leading to neurological conditions such as pain. Indeed, a recent study looked at differential gene expression in DRGs taken from both nerve injury and inflammation-induced mouse pain models and identified the ‘ECM organization’ pathway as being the most dysregulated [57]. In addition, another study found that a subset of collagens (*COL3A1, COL9A2, COL6A2*, and *COL5A1*) were upregulated in the DRGs of male individuals with neuropathic pain compared to pain-free controls [33]. Clinically, there are also examples of collagen-related diseases with unexplained neurological effects such as pain [9]. For example, several heritable connective tissue disorders caused by defects in collagen-encoding or collagen-regulating genes display a pain phenotype, which is phenocopied in their corresponding murine models (e.g., *Col1a1*^*Jrt/+*^ and *Col5a1*^*+/-*^ mice), suggesting that healthy core matrisome and matrisome-associated signaling is crucial for normal neuronal function and pain perception [58,59]. Receptors that can bind collagen have also been shown to modulate pain. In the literature, CD44 binding with its other ligand, hyaluronic acid, has been shown to have anti-hyperalgesic effects [60]. In contrast, integrin signaling has been implicated in maintaining hyperalgesia in both neuropathic and inflammatory rat models of pain [61,62]. In both cases, the role of collagen interactions with these receptors in painful conditions has not been investigated. Finally, fibroblasts in the DRG have also recently been shown to produce a protein, secreted modular calcium-binding protein 2, that may play a role in suppressing mechanical allodynia by inhibiting neuron-neuron interactions in the DRG [63]. Taken together, it can be appreciated that the ECM plays an important role in the physiological functioning of the peripheral nervous system.

This study has a few limitations. First, while we have similar numbers per sex for the age matched murine samples, a limitation of this study is that the human bulk RNAseq samples have more variability in age and numbers per group with 11 male and 4 female samples. Therefore, we did not directly assess sex differences. In addition, a more age-matched approach would be preferred but is challenging given the limited availability of human donor samples. Secondly for our scRNAseq experiment, we only used male mice, which inhibited our ability to look for sex-specific differences at this level. Finally, one should note that this study and its calculations are based mainly on transcriptomics data, which does not always correlate to the protein level and the downstream effects of protein activities.

Overall, this work adds to the literature by establishing the prevalence of matrisome gene expression in the DRG, identifying the cellular components producing these molecules, and providing insight on collagens and their interaction partners in mouse and human DRG. This study will serve as a framework for future examination of the functional consequences of these matrisome expression patterns. It will be interesting to further investigate how the ECM interactions with neurons potentially contribute to pain as well as to other neurological conditions.

## Experimental procedures

### Murine bulk RNA sequencing

Animal experiments were approved by the Ethical Committee of Ghent University (ECD20-62). Mice were housed 2 to 5 per cage with food and water *ad libitum* and kept on 12-hour light cycles. 15-week-old male (n = 6) and female (n = 5) wild-type C57BL/6 mice were euthanized with CO2 asphyxiation and bilateral lumbar L3-L5 DRG were collected under RNase free conditions, snap frozen and stored at −80°C. Subsequently, RNA extraction was performed using the RNeasy kit with on column DNase digestion as recommended by the manufacturer (Qiagen). Bulk RNA sequencing was performed using TruSeq Stranded mRNA library prep followed by 150bp paired-end sequencing on Illumina’s NovaSeq6000 to obtain 30 million paired-end reads per sample. Reads were aligned against the mouse reference genome (GRCm38) with STAR and counted with StringTie v2.0. TPM values were calculated. To determine the cutoff TPM value above which genes were considered expressed, the average TPM value was calculated for all matrisome genes that were only expressed by one of the 11 samples. This led to a cutoff TPM value of 0.1. A publicly available *in silico* list of murine matrisome genes (n = 1,110, v2.0 http://matrisomeproject.mit.edu/other-resources/mouse-matrisome/) was used to filter the bulk RNA sequencing data. For 12 genes listed in the murine matrisome list, the provided gene name was not found in the bulk RNAseq data. For six genes an alias was found that allowed detection in the bulk RNAseq dataset when replaced in the murine matrisome gene list. For six genes (*Ntn3, Itlnb, Lgals6, Gm5347, U06147, Prl2c4*), no alias could be identified and no match in the murine bulk RNAseq data was found.

### Human DRG bulk RNA sequencing

Previously published human DRG bulk RNAseq data of 15 donors (male: n = 11, female: n = 4, minimum age: 37, median: 61, maximum: 79 years) was provided by the lab of Dr. Theodore J. Price and can be found on the website (https://paincenter.utdallas.edu/sensoryomics/) [33]. Patients that were classified with no neuropathic pain were selected for this study. A predefined *in silico* list of human matrisome genes (n = 1027, v2.0 http://matrisomeproject.mit.edu/other-resources/human-matrisome/) was used to filter the human bulk RNA sequencing data. TPM values were calculated and an average TPM value above 0.9 was considered expressed, by averaging the TPM values of all genes that were expressed in only one of the 15 samples. For 20 genes in the human matrisome list, no match was found for the provided gene name in the human bulk RNA sequencing dataset. Upon checking the HUGO Gene Nomenclature database (https://www.genenames.org) an alias or approved gene name could be identified for 14 of these genes and replacing the listed gene name with the alias allowed detection in the bulk RNAseq dataset. For the six remaining genes (*MUC19, MUC2, MUC8, SERPINA2, CCL4L1, MST1L*), no alias could be identified, hence no match could be found in the human bulk RNAseq data.

### Murine single cell RNA sequencing

This experiment was approved by the Institutional Animal Care and Use Committees at Rush University Medical Center and Northwestern University. Animals were housed with food and water *ad libitum* and kept on 12-hour light cycles. Single cell RNA sequencing (scRNAseq) was performed on pooled L3-L5 DRG cells collected unilaterally from 10, 18-week-old male C57BL/6 mice as described [64]. Cells were dissociated and cell number and viability were analyzed using Nexcelom Cellometer Auto2000 with AOPI fluorescent staining method. Single cell gel beads were generated using 10x Genomics Chromium controller chips at the Northwestern University sequencing core. cDNA and library preparation were performed using 10X Genomics Chromium kits, and samples were sequenced using 50bp paired-end HiSeq sequencing. Sequencing reads from 9,400 cells were assembled and aligned against the mouse reference genome using the 10x Genomics Cell Ranger v6.0.0. Expression count matrices were analyzed using the Seurat (v4.0.1) R package. Downstream analysis was performed as described before resulting in 8,755 cells for analysis [64]. Cluster names were determined by comparing expression profiles of markers per cluster with mousebrain.org and celltypist.org databases (Supplemental Figure S3A-B) [34–36]. Raw fastq files and the expression count matrix have been deposited on NCBI GEO (accession number GSE198485).

### Intercellular communication analysis

Metadata and data slots of the Seurat object were used to generate a CellChat object using the CellChat R package (CellChat 1.1.3) [38]. The murine DRG scRNAseq data was preprocessed using CellChat’s standard workflow. CellChat’s database of 2,021 known ligand-receptor interactions in mice (http://www.cellchat.org/cellchatdb/) was used to infer ligand-receptor interactions and standard pre-processing functions of *identifyOverExpressedGenes* and *identifyOverExpressedInteractions* were applied with CellChat’s default parameters. Cell-cell communication probability was calculated and communications with fewer than 10 participating cells excluded from analysis. Here we focused on ECM related pathway networks. Aggregated cell-cell communication as well as cell-cell communication for the signaling pathways of interest were calculated. Chord, circle, and hierarchy plots were generated using the *netVisual_aggregate()* and *netAnalysis_contribution()* functions.

### Human DRG spatial transcriptomics

Previously published human DRG Visium spatial transcriptomics data of 8 donors (Male: n = 4, female: n = 4) was provided by the lab of Dr. Theodore J. Price and can be found on the website (https://paincenter.utdallas.edu/sensoryomics/) [39]. For our in-house sample, one human DRG sample (female; age 96 years, BMI 22.5) was acquired from the ROS/MAP studies as described below (Rush University Medical Center). For both sets of samples, Visium tissue optimization and spatial gene expression protocols were followed as described in the manufacturers manual (https://10xgenomics.com/). Hematoxylin and eosin (H&E) was used as a counterstain for both. Imaging was conducted on an Olympus vs120 slide scanner (Price lab, UTDallas) or ZEISS LSM 980 with Airyscan 2 (Rush University Medical Center). mRNA library preparation and sequencing were done at the Genome Center in the University of Texas at Dallas Research Core Facilities (Illumina Nextseq 500) as previously published [39], or at Roy J. Carver Biotechnology Center at University of Illinois at Urbana-Champaign (Illumina NovaSeq 6000) for the in-house sample. The 10x Visium Spatial Tissue Optimization workflow was used to optimize permeabilization conditions for the in-house sample, and the optimal permeabilization time was determined to be 6 min. The data associated with the in-house sample can be found at NCBI GEO (accession number GSE215994).

To assign Visium barcodes to a certain cell type, *DCN* and *SCN10A* were selected as markers to identify VLMC-like and nociceptor barcodes, respectively. The 25^th^ quartile was calculated for all of the barcodes having scaled and normalized expression of the gene of interest above 0. Barcodes with expression above this 25^th^ quartile value were considered positive for the gene of interest. These calculations were repeated for selected collagens: *COL1A1, COL1A2, COL4A1*, COL4A2, *COL6A1, COL6A2* and predicted receptors: *CD44, SDC4, ITGA1, ITGB1, ITGA3, ITGAV*, and *ITGB8*. Subsequently the sum of double positive barcodes for the gene of interest and *SCN10A* were divided by the total number of *SCN10A* positive barcodes. In addition, the sum of double positive barcodes for the gene of interest and *DCN* were divided by the total number of *DCN* positive barcodes. Double positive barcodes for *SCN10A* and *DCN* were excluded. Finally, we compared the values of both ratios by a paired *t*-test for each of the 9 samples.

### Human DRG RNAscope

In house human DRG came from participants in the Religious Orders Study (ROS) or Rush Memory and Aging Project (MAP) [65]. At enrollment, participants agreed to annual clinical evaluation and organ donation at death, including brain, spinal cord, nerve, and muscle. Both studies were approved by an Institutional Review Board at Rush University Medical Center. All participants signed an informed consent, Anatomic Gift Act, and a repository consent to allow their resources to be shared. The DRGs were removed postmortem within 12 hours and flash frozen as part of the spinal cord removal. Two male human DRG samples (donor 1: age = 82.02 years, BMI = 21.26; donor 2: age = 95.2years, BMI = 25.17) were acquired from the ROS/MAP studies (Rush University Medical Center). ROSMAP resources can be requested at https://www.radc.rush.edu.

RNA *in situ* hybridization (ISH) was performed using ACD Bio-Techne RNAscope Multiplex Fluorescent v2 Assay. For human DRGs, modifications were made to the protocol to preserve tissue integrity. Briefly summarized, slides were removed from −80°C and immediately submerged in 4% PFA on ice for 40 min. Dehydration was performed following 50%, 75% and two 100% ethanol washes for 5 min each. Hydrogen peroxide (3%) was applied for 10 min. Target retrieval was performed, reducing time in target retrieval buffer to 3 min followed by protease III incubation for 30 min. The remainder of the protocol was performed following manufacturer’s instructions. Probes were used at 1:50 dilution and Opal dyes from Akoya Biosciences were used at 1:100 dilution. Opal dyes 570 (OP-001003) and 650 (OP-001005) were used. *SCN10A* (406291-C3), *COL1A1* (401891-C2), *COL3A1* (549431-C1) and *COL5A1* (1148401-C1) probes were used. For DAPI staining, Vectashield containing DAPI was used. ACD Bio-Techne positive and negative control probes were conducted prior to start of work. Negative controls were included on every slide. Imaging was performed using an Olympus Fluoview FV10i confocal microscope at 10x and 60x magnification. Multiple planes of focus were captured, but Z-stacks were not produced and instead the optimally focused image was chosen for processing and analysis. Laser intensity was used at ≤9.9% throughout. Images were processed and quantified using Fiji software. Only brightness and contrast tools were used to adjust images. In order to quantify cellular expression of *COL1A1, SCN10A* and *DCN*, 10 images at 60X magnification per human DRG were analyzed. First, the total number of cells was identified using both the nuclei staining with DAPI and the phase contrast channels. Each cell was then assessed for expression of each probe, and labeled as single expression, double expression, or no expression. Positive signal was determined when 2 or more positive ‘dots’ per cell were found. The total number of cells assessed is indicated in figures.

### Immunohistochemistry on human DRG

All human tissue procurement procedures were approved by the Institutional Review Boards at the University of Texas at Dallas. Human lumbar DRGs from one male and one female organ donor with no notable chronic pain conditions (21 and 58 years old, respectively) were collected within 4 hours of cross-clamp, frozen on dry ice and stored in a −80°C freezer until use. One L4 DRG from each donor was embedded in OCT and cut on a cryostat into 20µm sections that were applied directly onto SuperFrost Plus charged slides. Slides were submerged in 4°C 1% formalin for 15 minutes, then dehydrated in 50%, 70% and 100% ethanol for 5 minutes each. The slides were allowed to dry briefly then a boundary was drawn around the sections using a hydrophobic pen (ImmEdge PAP pen, Vector Labs) and placed in a light-protected, humidity-controlled tray. Sections were incubated overnight in rat anti-CD140a (PDGFRA) (ThermoFisher; #14-1401-82; RRID: AB_467491; 1:50) and rabbit anti-TRPV1 (ThermoFisher; #PA1-748; RRID: AB_2209010; 1:500) diluted in 0.1M phosphate buffer (PB), with 5% normal goat serum and 0.3% Triton-X 100 (PBS-T). Following this, sections were rinsed twice with PB then incubated with species-specific secondary antibodies diluted in PBS-T for two hours (goat anti-rat conjugated to Alexa 488; ThermoFisher; #A-11006; 1:500 and goat anti-rabbit conjugated to Alexa 647; ThermoFisher; A-21247; 1:500). After two final rinses in PB, a cover slip with a small volume of mounting medium (4% n-propyl gallate, 85% glycerol and 10 mM phosphate buffer, pH 7.4) was applied and secured using nail varnish. Sections were scanned using an Olympus FV3000RS confocal microscope with a x40 oil-immersion lens and a 1.5x zoom. During image acquisition, an empty channel was scanned to visualize autofluorescence, including lipofuscin.

### Statistical analyses

All analyses were carried out in Microsoft Excel, Graphpad Prism 9.4.0 (GraphPad Software, San Diego, CA), the Seurat (v4.0.1) R package, or the Cell Chat (v1.1.3) R package. Bulk RNAseq averages and cutoffs were calculated and ranked in Excel and graphs were created with Graphpad Prism. Core matrisome versus matrisome-associated gene comparisons using bulk RNAseq data were analyzed with unpaired two-tailed t-test. Human spatial transcriptomics data were analyzed in Seurat and co-expression with cellular markers was compared with a paired two-tailed *t*-test. Data are expressed as mean±SEM, with *n* indicating the number of samples. P values less than 0.05 were considered significant.

## Author contributions

**Robin Vroman**: Conceptualization, study design, software, data analysis, interpretation of results, writing of the manuscript, **Rahel Hunter**: software, data analysis, reviewing and editing of the manuscript, **Matthew J. Wood**: data acquisition, data analysis, reviewing and editing of the manuscript, **Olivia C. Davis:** data acquisition, data analysis, reviewing and editing of the manuscript, **Zoë Malfait**: data acquisition, data analysis, reviewing and editing of the manuscript, **Dale S. George:** data acquisition, reviewing and editing of the manuscript, **Dongjun Ren:** data acquisition, reviewing and editing of the manuscript, **Diana Tavares-Ferreira:** data acquisition, reviewing and editing of the manuscript, **Theodore J. Price:** interpretation of results, reviewing and editing of the manuscript, **Anne-Marie Malfait**: conceptualization, interpretation of results, reviewing and editing of the manuscript, **Fransiska Malfait**: conceptualization, interpretation of results, reviewing and editing of the manuscript, **Rachel E. Miller**: conceptualization, study design, software, data acquisition, data analysis, interpretation of results, writing of the manuscript, reviewing and editing of the manuscript, **Delfien Syx**: conceptualization, study design, software, data acquisition, data analysis, interpretation of results, writing of the manuscript, reviewing and editing of the manuscript.

## Acknowledgements

The Northwestern University NUSeq Core Facility is acknowledged for performing the single cell RNAseq work. The 10x Genomics Chromium System employed for the scRNA-seq is made available with an NIH S10 Grant to NUSeq (1S10OD025120). The Carver Biotechnology Center at the University of Illinois Urbana-Champaign is acknowledged for performing the Visium spatial transcriptomics experiment of the in-house sample. We thank the study participants and staff of the Rush Alzheimer’s Disease Center. ROSMAP resources can be requested at https://www.radc.rush.edu.

## Abbreviations

*Adam*: a disintegrin and metalloproteinase domain
*Anxa*: annexin
*Cd44*: cluster of differentiation 44
*Col*: collagen
*Cts*: cathepsin
DAPI: 4ʹ,6-diamidino-2-phenylindole
DCN: decorin
DRG: dorsal root ganglion
ECM: extracellular matrix
*Fgf*: fibroblast growth factor
FISH: fluorescence *in situ* hybridization
*Fn1*: fibronectin 1
H&E: hematoxylin and eosin
*Hspg*: heparan sulfate proteoglycan
IHC: immunohistochemistry
IMM: immune cells
*Itg*: integrin
*Lam*: laminin
LDN: large diameter neurons
*Lgals*: galectin
MAP: memory and aging project
NOCI: nociceptor
*Ogn*: osteoglycin
PDGFRA: platelet derived growth factor receptor α
PFA: paraformaldehyde
RNAseq: RNA sequencing
ROS: religious orders study
SATG: satellite glial cell
SCHW: Schwann cell
*SCN10A*: voltage-gated sodium channel alpha subunit 10
scRNAseq: single cell RNA sequencing
*Sdc*: syndecan
SEM: standard error of the mean
*Sema*: semaphorin
*Sfrp*: secreted frizzled related protein
*Srgn*: serglycin;
TPM: transcripts per million
*Trpa1*: transient receptor potential cation channel, subfamily A, member 1
*Trpv1*: transient receptor potential cation channel, subfamily V, member 1
*VEC*: vascular endothelial cell
*Vegf*: vascular endothelial growth factor
VLMC-like: vascular leptomeningeal-like cells
*VSMCA*: vascular smooth muscle cells, arterial

