## Supplemental Figures for "The matrisome of the murine and human dorsal root ganglion: a transcriptomal approach"

### Figure S1

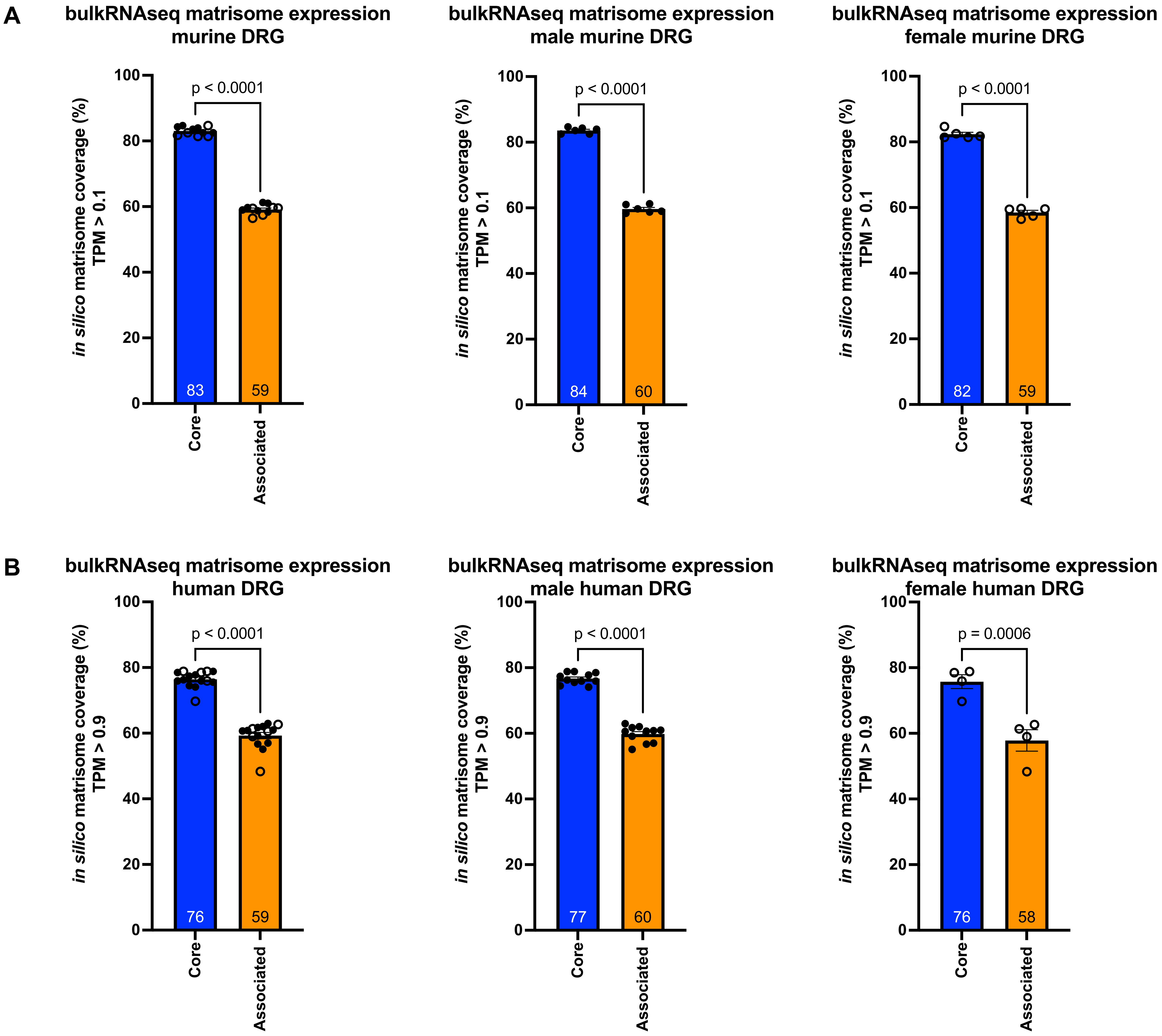


**Figure S1**: Matrisome expression in murine and human DRG. **A)** Percentage of expressed genes for core matrisome genes (total 274) vs matrisome associated genes (total 836) calculated from bulk RNAseq of murine DRG. From left to right, male and female plotted together or separately (n = 6 male (filled dots), n = 5 female (open dots), paired *t*-test); **B)** Percentage of expressed genes for core matrisome genes (total 274) vs matrisome associated genes (total 753) calculated from bulk RNAseq of matrisome genes in human DRG. From left to right, male and female plotted together or separately (n = 11 male (filled dots), n = 4 female (open dots), paired *t*-test). Number inside the bar represent the mean per group. Mean±SEM. Human bulk RNAseq data was previously published [33].

### Figure S2

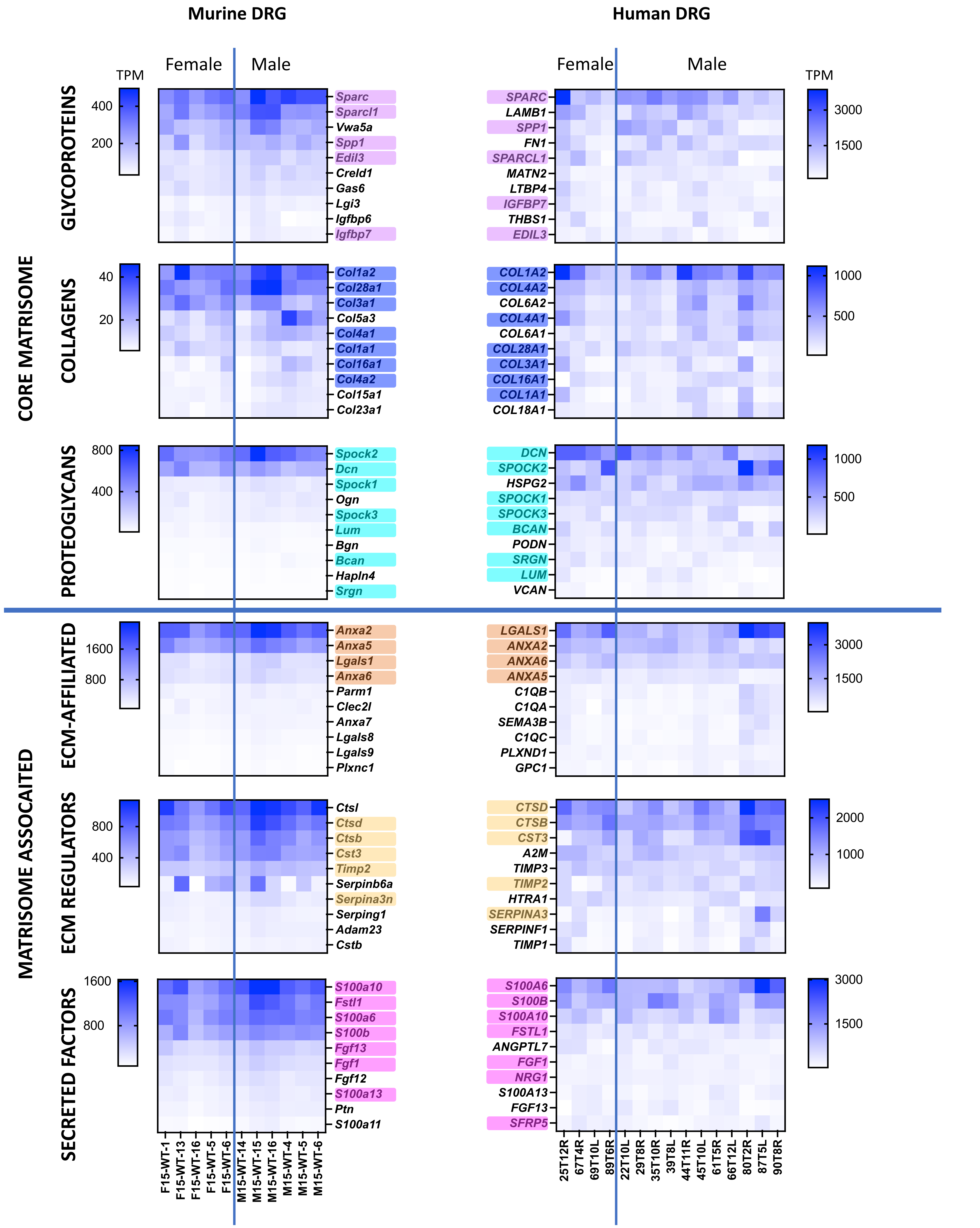


**Figure S2**: Overlap of 10 highest expressed matrisome genes between murine and human DRG per matrisome category based on bulk RNAseq. Highest expressed genes are ranked by average TPM value across all samples per matrisome category. (murine n = 6 male, n = 5 female; human n = 11 male, n = 4 female) Human bulk RNAseq data was previously published [33].

### Figure S3

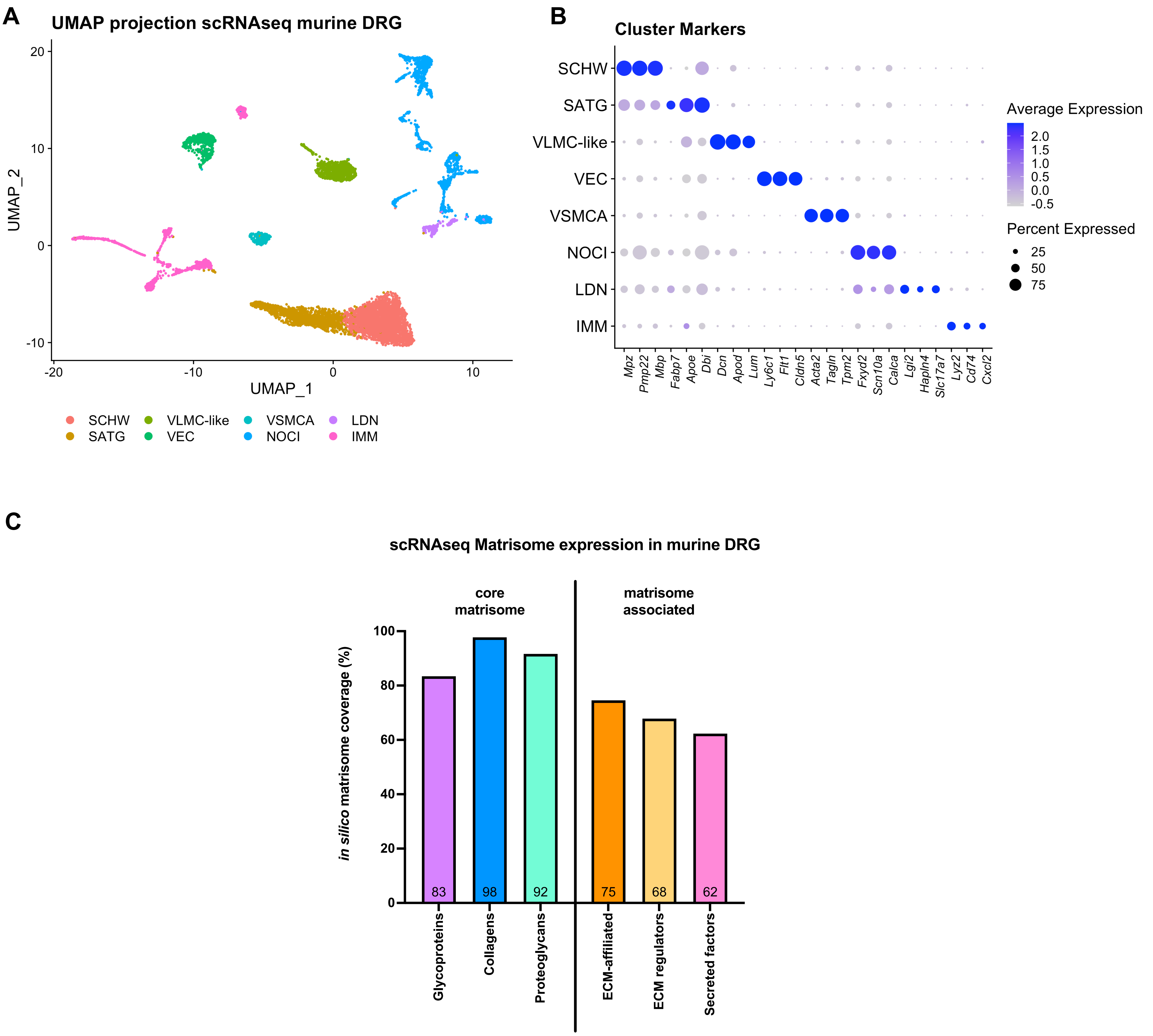


D

| | Total | SCHW | SATG | VLMC-like | VEC | VSMCA | NOCI | LDN | IMM |
|---|---|---|---|---|---|---|---|---|---|
| **number** | 8755 | 2801 | 1483 | 790 | 587 | 265 | 1614 | 178 | 1037 |
| **percentage** | 100 | 31.9 | 16.9 | 9.02 | 6.70 | 3.03 | 18.4 | 2.03 | 11.8 |

**Figure S3**: ScRNAseq on murine DRG. **A)** UMAP projection of scRNAseq of murine DRG, pooled unilateral L3-L5 DRG of 10 male mice at the age of 18 weeks. Different clusters were identified: Schwann cells (SCHW), satellite glial cells (SATG), vascular leptomeningeal-/fibroblast- like cells (VLMC-like), vascular endothelial cells (VEC), vascular smooth muscle cells arterial (VSMCA), nociceptors (NOCI), large diameter neurons (LDN), and immune cells (IMM). **B)** Clusters were defined using markers identified with the FindMarkers command in Seurat. The size of the dot represents the percentage of cell expressing the given gene within a cluster, and the color corresponds to the average expression (unscaled data) across all cells within a cluster for each gene of interest. **C)** Matrisome genes expression in murine DRG bulkRNAseq data, bar chart of percentage of expressed genes for each category of the matrisome. Number inside the bar represent the mean per group. **D)** Numbers and percentages of cells in each cluster.

### Figure S4

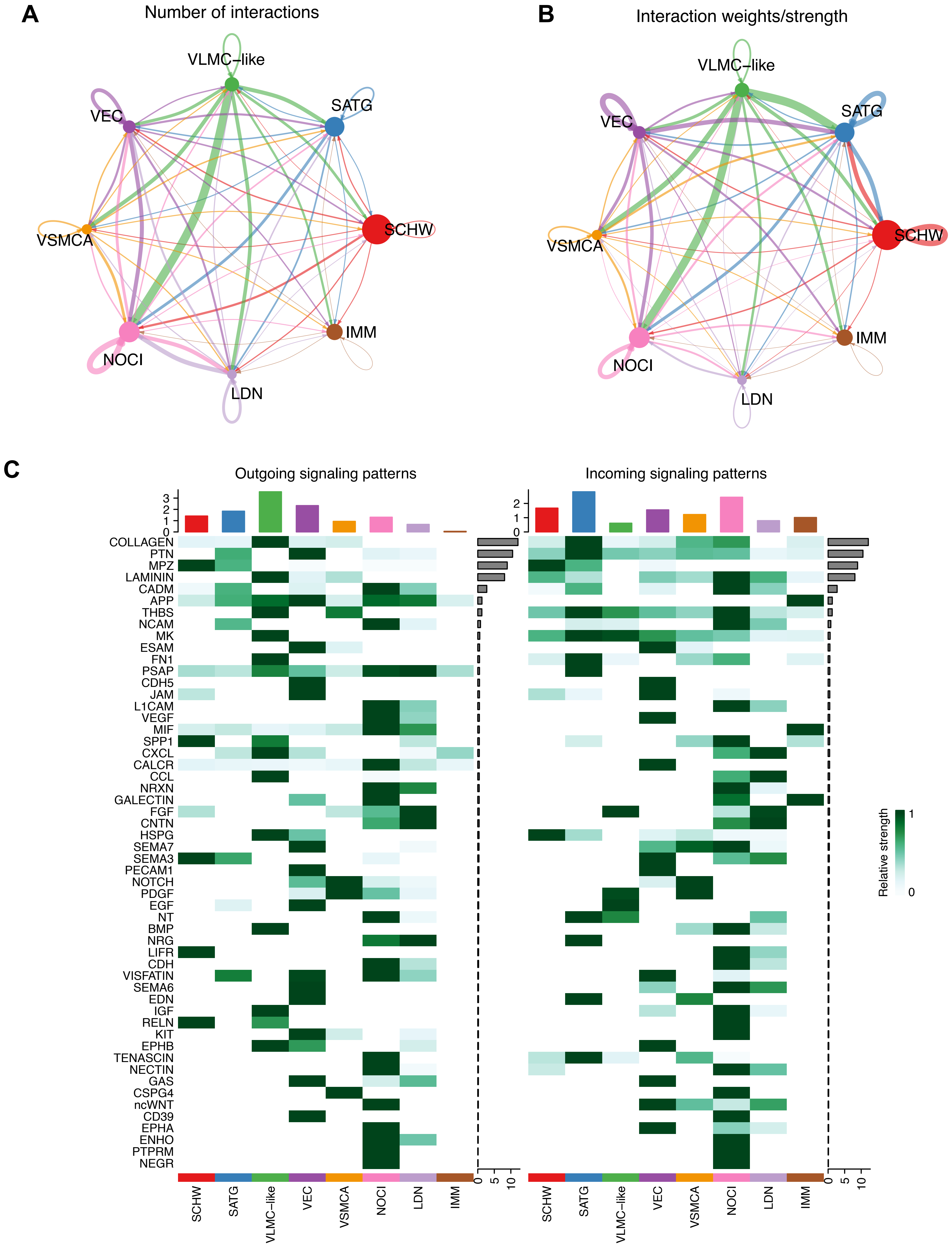


**Figure S4:** Cell chat inferred cell-cell communication and pathway analysis. **A)** Absolute number of inferred interactions in the murine DRG tissue. Arrowhead show the direction of interaction while the thickness of the arrow is representative for the number of interactions. **B)** Inferred interactions strength (weights) are represented. Arrowhead show the direction of interaction while the thickness of the arrow is representative for the weight/strength of the interaction. Size of the dot representing the cell type correlates with the number of cells for that cell type. **C)** Outgoing and incoming signaling role analysis on the aggregated cell-cell communication network from all found pathways. All ligand-receptor pathways are ranked based on their weight in the DRG sample. Outgoing signaling patterns show the cell type sources of the ligands of the interactions of the pathway while incoming signaling patterns indicate which cell types express the receptors of the pathway. The height of the colored bar chart on top represents the total strength of each cell type of as a source of all aggregated interactions, and the height of the right grey bar indicates the strength of the signaling pathway by summarizing all cell types. Schwann cells (SCHW), satellite glial cells (SATG), vascular leptomeningeal-/fibroblast- like cells (VLMC-like), vascular endothelial cells (VEC), vascular smooth muscle cells arterial (VSMCA), nociceptors (NOCI), large diameter neurons (LDN), and immune cells (IMM).

### Figure S5

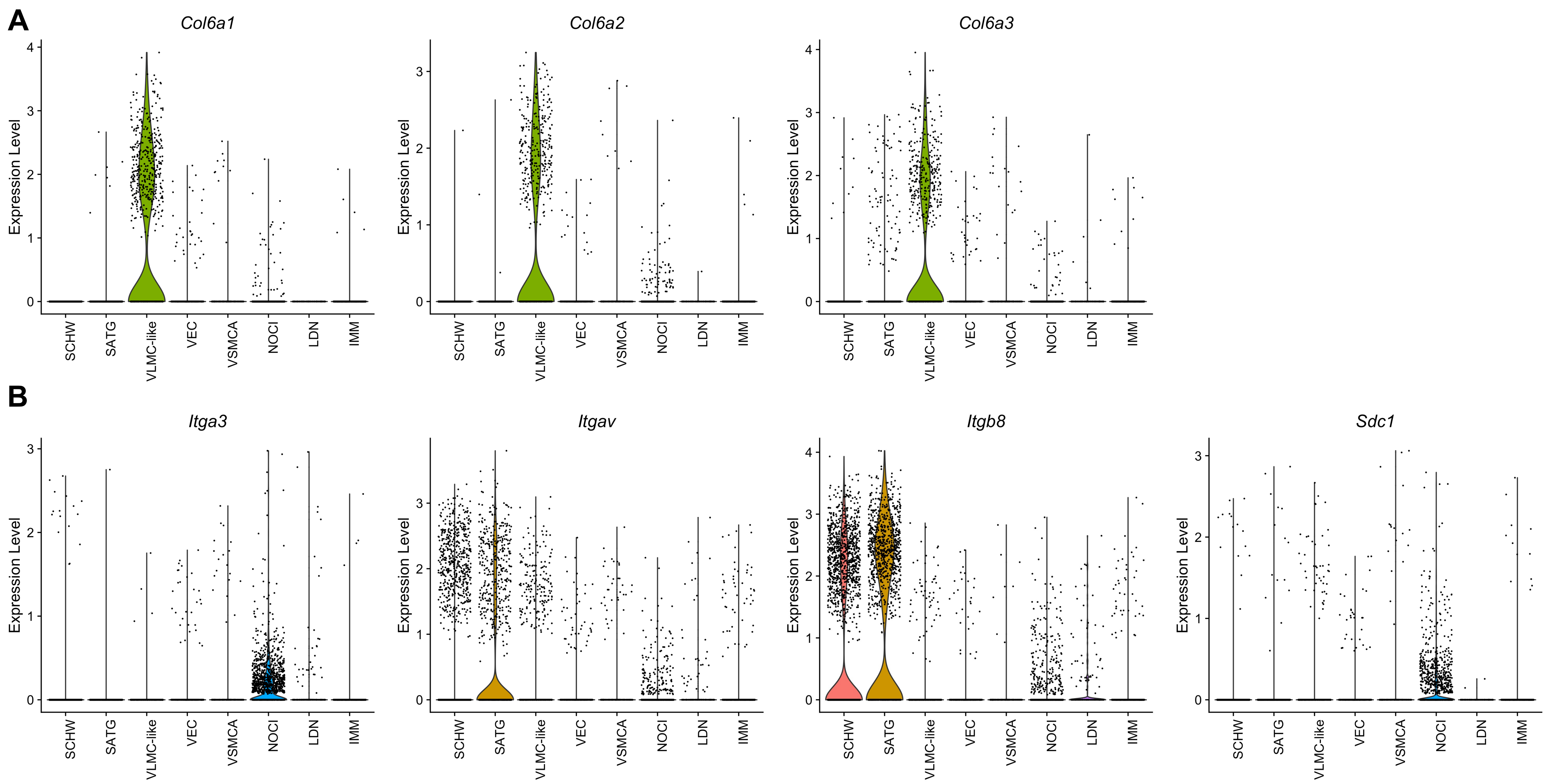


**Figure S5**. Violin plots of other highly contributing ligand-receptor pairs in the collagen pathway network in scRNAseq murine DRG associated with Figure 4B. **A)** The ligands and their expression profiles over the different clusters. **B)** The receptors and their expression profiles over the different clusters. Schwann cells (SCHW), satellite glial cells (SATG), vascular leptomeningeal-/fibroblast- like cells (VLMC-like), vascular endothelial cells (VEC), vascular smooth muscle cells arterial (VSMCA), nociceptors (NOCI), large diameter neurons (LDN), and immune cells (IMM).

### Figure S6

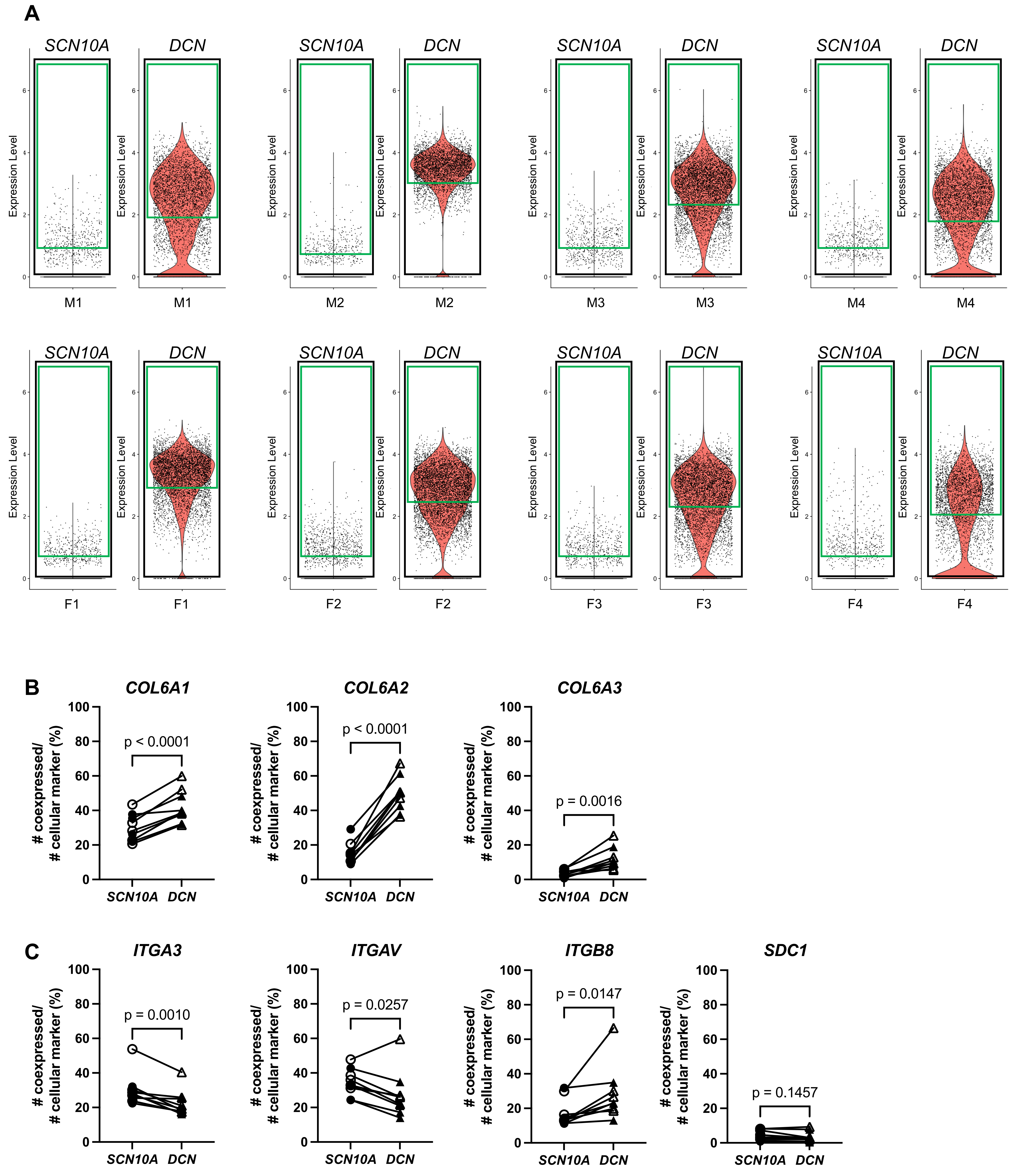


**Figure S6**: Spatial transcriptomics on human DRG. **A)** After scaling and normalization in Seurat, expression thresholds were established for each gene of interest by using the 25th quartile cutoff (green box) of barcodes > 0 (black box) for that gene in each DRG sample. *SCN10A* and *DCN* are shown here as examples for one DRG, but this method was also applied to genes in parts B and C. **B)** Percentage of co-expression of *COL6A1, COL6A2 and COL6A3* respectively with *SCN10A* or *DCN*. **C)** Percentage of co-expression of *ITGA3, ITGAV* and *ITGB8* respectively with *SCN10A* or *DCN. SCN10A-DCN* double positive cells were excluded from analyses. n = 4 male (filled symbol), n = 5 female (open symbol).

### Figure S7

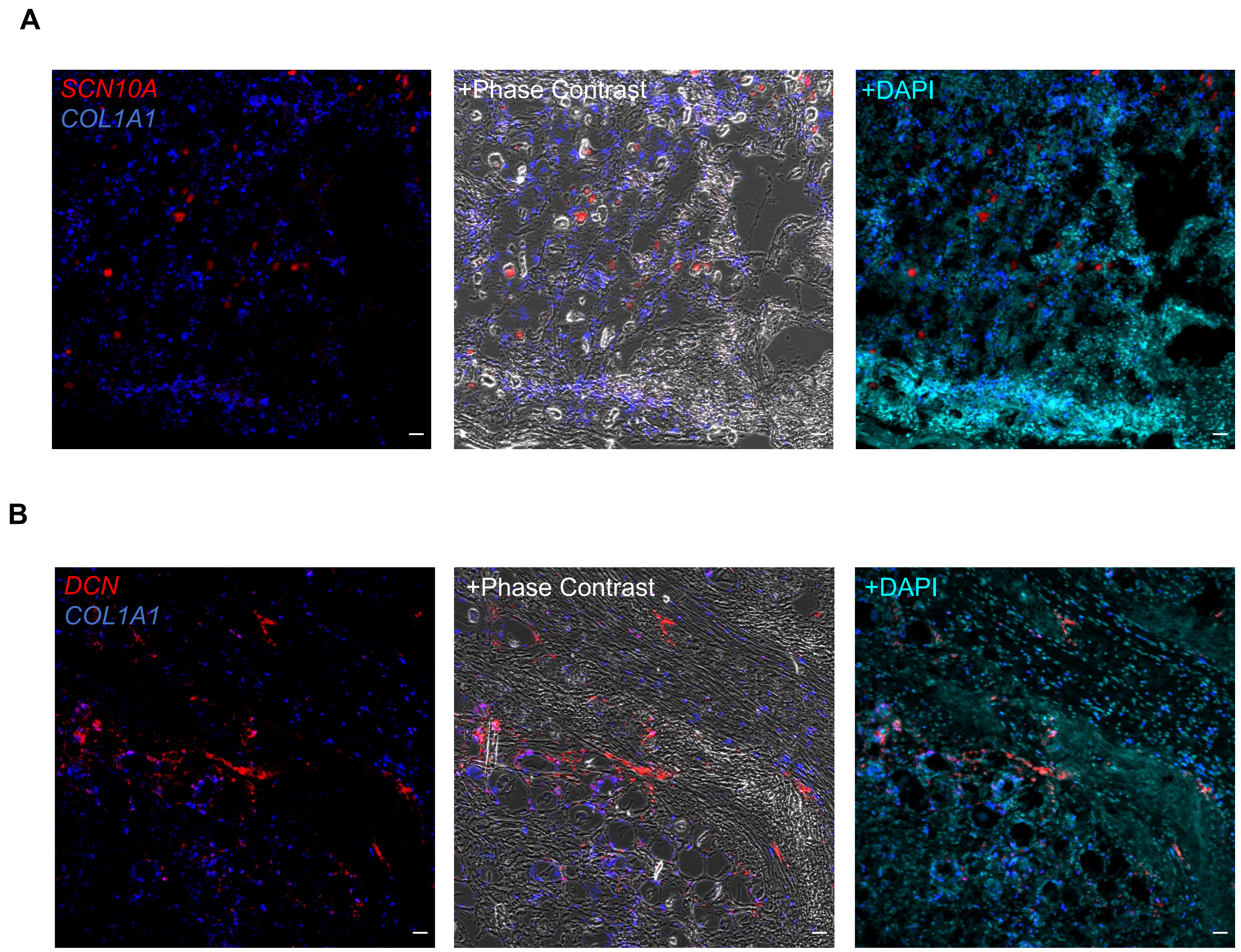


**Figure S7:** Expression of *COL1A1, SCN10A* and *DCN* in Human DRG. **A)** RNAscope used to identify cells expressing *SCN10A* and *COL1A1* or **B)** *DCN* and *COL1A1* in human DRG (male donors, n=2). Representative sections are shown for *COL1A1* staining with the cellular marker, either with phase contrast or nuclei staining (DAPI) overlay. Scale bar = 50µm (A,B).
